# Structural Basis for Substrate Binding, Catalysis and Inhibition of Breast Cancer Target Mitochondrial Creatine Kinase by Covalent Inhibitor via Cryo-EM

**DOI:** 10.1101/2024.06.18.598884

**Authors:** Merve Demir, Laura Koepping, Ya Li, Lynn Fujimoto, Andrey Bobkov, Jianhua Zhao, Taro Hitosugi, Eduard Sergienko

## Abstract

Mitochondrial creatine kinases are key players in maintaining energy homeostasis in cells by working in conjunction with cytosolic creatine kinases for energy transport from mitochondria to cytoplasm. High levels of MtCK observed in Her2+ breast cancer and inhibition of breast cancer cell growth by substrate analog, cyclocreatine, indicate dependence of cancer cells on the ‘energy shuttle’ for cell growth and survival. Hence, understanding the key mechanistic features of creatine kinases and their inhibition plays an important role in the development of cancer therapeutics. Herein, we present the mutational and structural investigation on understudied ubiquitous mitochondrial creatine kinase (uMtCK). Our cryo-EM structures and biochemical data on uMtCK showed closure of the loop comprising residue His61 is specific to and relies on creatine binding and the reaction mechanism of phosphoryl transfer depends on electrostatics in the active site. In addition, the previously identified covalent inhibitor CKi showed inhibition in breast cancer BT474 cells, however our biochemical and structural data indicated that CKi is not a potent inhibitor for breast cancer due to strong dependency on the covalent link formation and inability to induce conformational changes upon binding.

## Introduction

Creatine kinases (CKs) play a key role in energy homeostasis by facilitating energy transport from the mitochondria to the cytoplasm^1,2^. The ubiquitous and sarcomeric mitochondrial isoforms of CKs (uMtCK and sMtCK, respectively) are octameric enzymes residing in the intermembrane space of mitochondria that catalyze the reversible transfer of the phosphoryl group from ATP generated by mitochondria to creatine (Cr) to form ADP and phosphocreatine (pCr)^3–5^. Phosphocreatine allows accumulation of the high energy phosphate groups in the vicinity of cellular processes with high energy needs. Dimeric cytosolic creatine kinases (cCKs) work in conjunction with MtCK to convert pCr into ATP and restore the ATP/ADP levels in the cells for energy buffering (**Figure 1a**). The interplay between CKs enables cells to maintain the energy supply and homeostasis^6^. These functions are especially crucial in cancer cells to satisfy high energy demand for cell growth and survival^7^. Studies demonstrated increased level of uMtCK protein in Her2+ breast cancer and reliance of tumor cells on uMtCK activity^8,9^. Similar observations were made in EVI+ AML cells^10^. A creatine analog, cyclocreatine, has been shown to inhibit the growth of breast cancer that underlines the importance of understanding the mechanism of substrate binding and catalysis by uMtCK as a major step towards the development of cancer therapeutics^8^.

**Figure 1.**
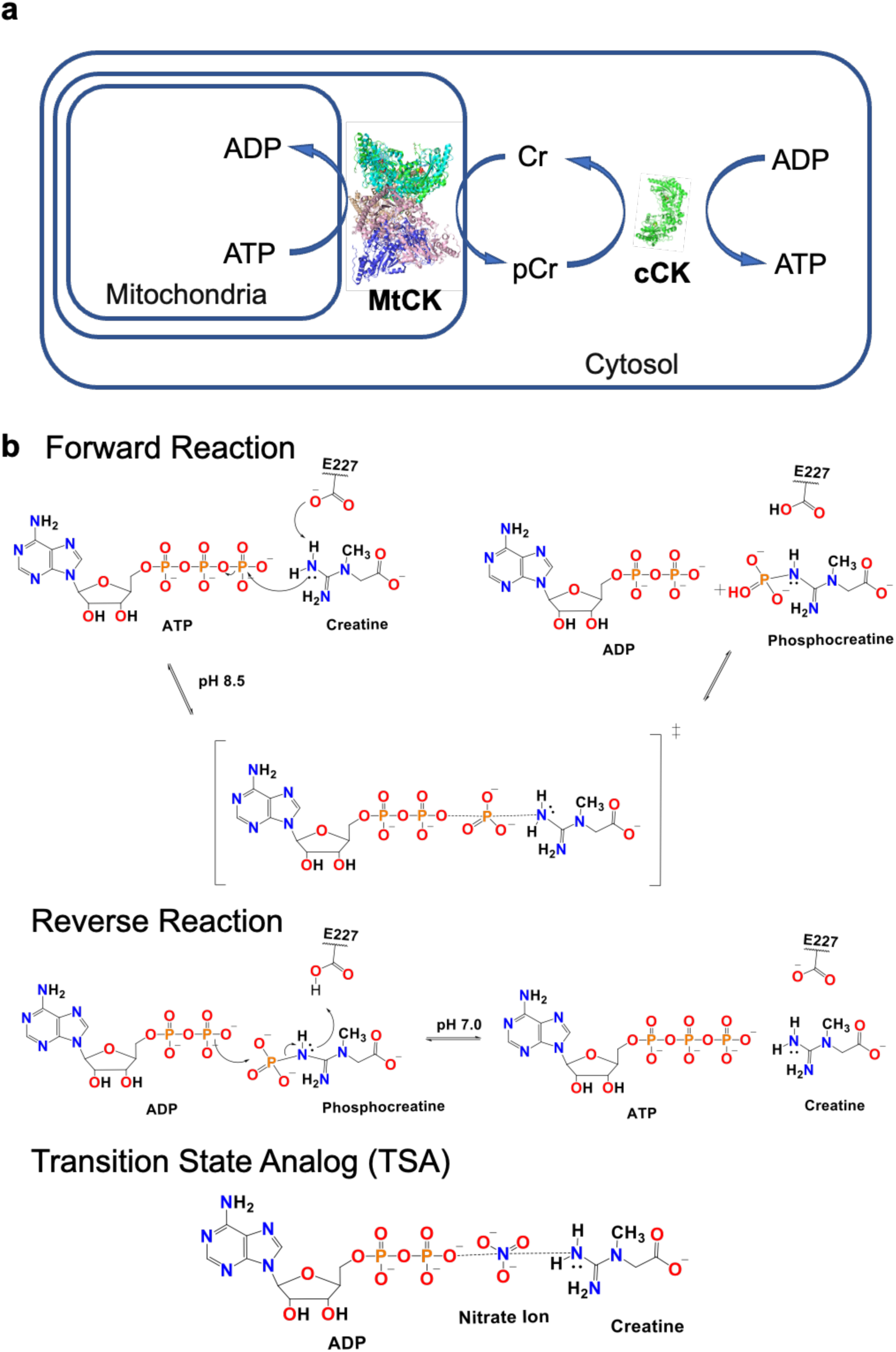
The interplay between CKs and phosphoryl transfer mechanism a) MtCK located in the intermembrane of mitochondria produces pCr that is used by cCKs in cytosol to generate ATP needed for cell function and energy homeostasis. b) Proposed forward and reverse reaction mechanisms of the phosphoryl transfer by CKs and transition state analog that mimics the transition state of the reaction are presented. Transition state analog has been used to obtain structures of CKs previously and in this study to understand the mechanism of the enzymes.

Previous studies on cCKs and uMtCK proposed an S_N_2 mechanism for the phosphoryl transfer with acid-base catalysis by Glu227 (residue numbers refer to the sequence of human ubiquitous MtCK protein without mitochondrial targeting sequence, residues 1-38)^11,12^. According to this mechanism, Glu227 acts as a general base in the forward reaction to remove a proton from the nucleophilic nitrogen of the guanidino group of creatine to enable the transfer of the phosphoryl group resulting in phosphocreatine (**Figure 1b**). The reversed reaction of phosphate transfer from phosphocreatine to ADP is accelerated at lower pH values where Glu227 acts as a general acid to protonate the nitrogen of creatine and phosphoryl group is transferred to ADP by a nucleophilic attack. Since the creatine kinase reaction does not generate stable intermediates, a transition state analog (TSA) comprised of ADP, Cr and NO_3_^-^ was used to mimic the step of the phosphoryl group transfer and to capture structures of cCKs that led to new findings on function of loops and residues involved in substrate binding and catalysis^13–17^. The mutational and structural studies of sMtCK and cCKs helped in understanding the roles for residues Glu227, Glu226, Asp321 and His61^18^.

Mutational studies on Glu227 and Glu226 showed their importance for catalytic activity, while loop residues Asp321 and His61, and active site residue Cys278 appeared to have a role in substrate binding^19–23^. In the apo-form of cCK, the active site is widely open and loops (referred to as Loop 1 and Loop 2 in this work) containing the aforementioned residues are disordered or not visible^17,24–26^. TSA-bound structures of cCKs displayed one monomer of the dimer bound to TSA with Loop 1 (comprising Asp321) and Loop 2 (containing H61) closing over the active site once the substrates are bound and Asp321 and His61 form a salt bridge, while the other monomer was found bound only to ADP with only the Loop 1 closed over the ADP binding site^14–16^.

The structural and mechanistic data for uMtCK1 are limited^11,27,28^. The existing X-ray crystal structure of human MtCK1 and Cryo-EM structure of bovine enzyme represent the protein in the absence of substrates (apo-MtCK1) where Loop 1 and Loop 2 were found in open conformations or missing density indicating disordered structure^29,30^. In this study, we present the first cryo-EM structures of human uMtCK in the presence or absence of substrates or TSA that reveal mode of substrate binding and concomitant conformational changes. We conducted mutagenesis of residues Glu227, Glu226, Asp321 and His61 to further our understanding of binding cooperativity and release of substrates as well as the reaction mechanism. Finally, we tested the previously discovered creatine kinase covalent inhibitor *in vitro* and in cellular assays to evaluate its effect on breast cancer^31^.

## Results and Discussion

### Structure Determination of Apo-, ADP- and TSA-MtCK

In our efforts to unravel the substrate binding and reaction mechanism of uMtCK, we determined the cryo-EM structure of WT human uMtCK in the presence and absence of substrates (**Figure 2**). The list of diverse MtCK1 cryo-EM structures solved in this study with the data processing and map statistics can be found in **Figure S1-2** and **Table S3**. The density map of WT uMtCK in the absence of substrates (apo-MtCK_WT_) with a resolution of 2.4 Å (**Figure 2a**) lacks the density for Loop 1 indicating its flexibility prevents it from being captured in a single stable conformation, while Loop 2 was found in an open conformation as also shown in previously reported structures of MtCK (**Figure 2c**)^29,30^.

**Figure 2.**
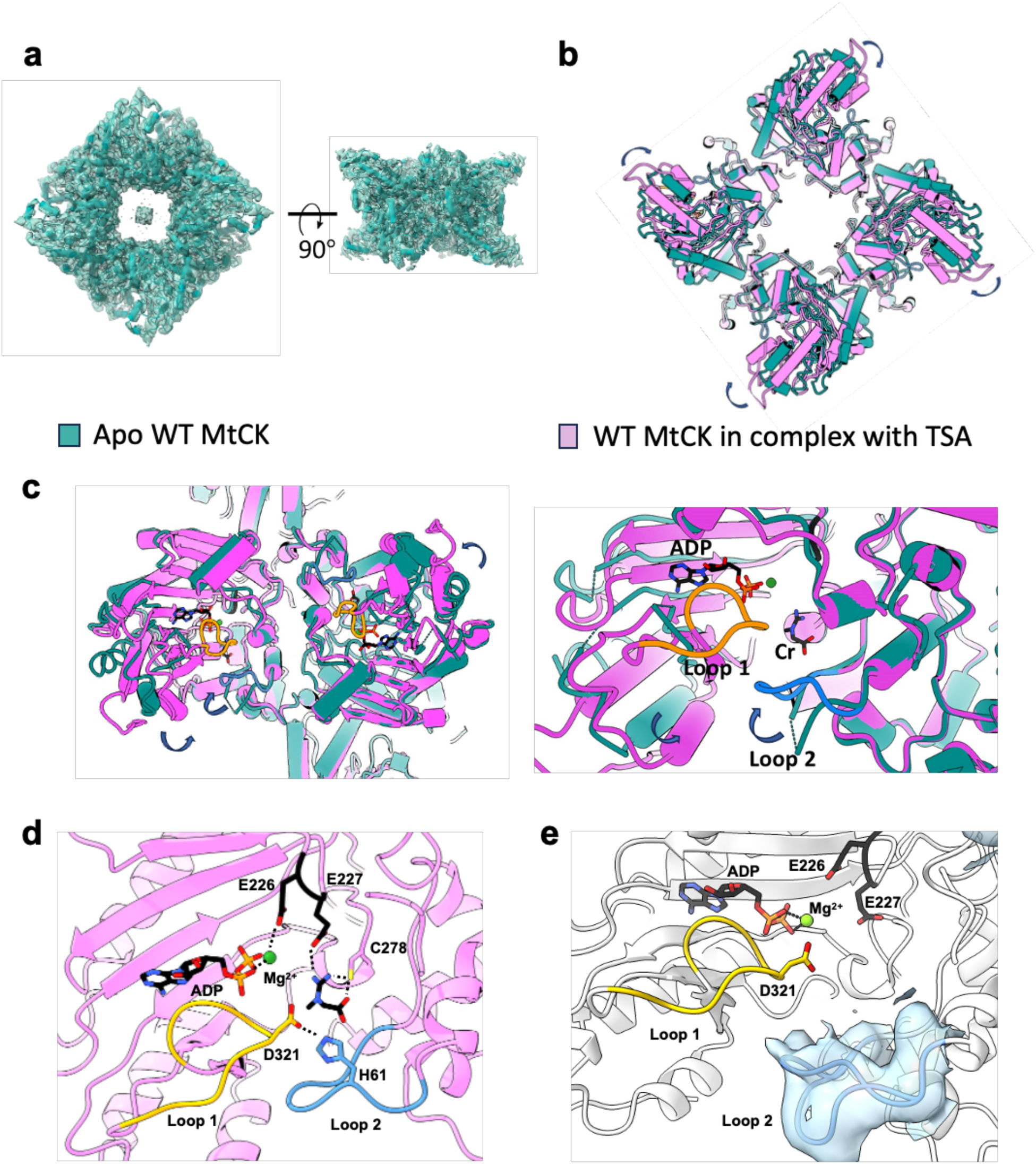
Cryo-EM structures of uMtCK a) The overall structure of apo-MtCK_WT_ determined using density map obtained with 2.4 Å resolution. b) Comparison of apo-MtCK_WT_ and TSA-MtCK_WT_ structures exhibit conformational changes from global movement of the C-terminal domains that **c)** contract and move in an inward motion upon closure of the flexible Loop 1 and 2 upon substrate binding. d) Active site structure of TSA-MtCK_WT_ highlights important residues Glu226, Glu227, Asp321 and His61 for ADP and Cr substrate binding. e) ADP-MtCK_WT_ structure shows ADP bound in the active site with Loop 1 closure over the substrate while Loop 2 takes a major closed and a minor open conformation as supported by the density map shown in blue.

The cryo-EM structures of uMtCK bound to ADP (ADP-MtCK_WT_) or a transition state analog (TSA-MTCK_WT_), were determined at 2.52 and 2.20 Å resolution, respectively. For obtaining the TSA-MTCK_WT_, while there are various ionic groups that can mimic the phosphoryl group that is transferred during the catalytic reaction^13^, we selected NO_3_^-^ to mimic the transfer of phosphoryl group from ATP to Cr as previously used in cCK structures. We observed several additional densities in the TSA-MtCK_WT_ map that were not present in the apo-MtCK map. An extra density at the ATP binding site in TSA-MtCK_WT_ indicated that ADP is bound in the active site with a Mg^2+^ ion coordinating to its phosphate backbone. A small density near Glu227 and Cys278 suggested the presence of Cr in the Cr binding site. Similar to what has been previously observed for cCK in complex with TSA ^14–17^, Loop 1 closes over the bound ADP (**Figure 2c**). Loop 2, that we observed in an open flexible state in apo-MtCK_WT_, was now found in a closed state latching together with Loop 1 in TSA-MtCK_WT_. Each monomer of uMtCK was found bound to ADP and Cr with Loop 1 and 2 latched together closing over the active site upon substrate binding. Unexpectedly, we do not observe density for the phosphoryl mimic NO_3_^-^, even though the crystal structures of cCK have shown a clear density for NO_3_^-^ ion. Despite the similarity of CK structures, the absence of NO_3_^-^ in the uMtCK structure may indicate a difference in reaction mechanism between MtCKs versus the cCKs.

Comparison of global changes in apo-MtCK_WT_ to TSA-MtCK_WT_ structures showed major movements for alpha helices in the C-terminal domain of the protein in addition to the loop movements upon substrate binding. The C-terminal alpha helices have contracted and move radially inwards, while the N-terminal remains largely unchanged (**Figure 2b-c**). Conformational changes in the ATP binding pocket allow insertion of adenine that forms interactions with the nearby histidine residues His186 and His291. These findings were further supported by the ADP-MtCK_WT_ structure where ADP is buried in the active site with Loop 1 closing over the substrate and Loop 2 adopting alternate conformations as evidenced by low occupancy of the closed conformation and high occupancy of the open conformation (**Figure 2e**). This indicates Cr is needed for loops to be latched together; however, the ability of Loop 2 to close in the presence of bound ADP is expected to lower the efficiency of Cr binding. In addition, we obtained the density map for uMtCK incubated with Cr alone which showed no substrate binding and was identical to apo-MtCK structure (data not shown). This result suggests Cr binding and loop closures are more effective in the presence of ATP and closure of Loop 1 may be required for Loop 2 closure. The overall conformational changes and movements of the alpha helices appear to aid the loops to latch together to catalyze transfer of the phosphoryl group, shield the active site from water access and avoid release of the substrates.

Upon comparison of cryo-EM structures we obtained, as well as with the previously published crystal structures, residues important for catalysis, substrate binding and order of loop closure were determined for mutational studies (**Figure 2d**). Residue Asp321 from Loop 1 interacts with His61 from Loop 2 that indicates a role for their involvement in forming the closed state of the loops upon substrate binding in a sequential manner. The residue Glu226 neighboring the catalytic Glu227 residue was shown to interact with Mg^2+^ ion that coordinates with the phosphate backbone of ADP. We set out to decipher the role of these residues in substrate binding, cooperativity and release by designing mutations that can impair their specific activity. In addition, we designed inactive mutants of uMtCK by replacing the catalytic residue, Glu227, to capture substrate- or product-bound structures of uMtCK that can help further our understanding of the reaction mechanism of uMtCK.

### Kinetic Characterization of uMtCK Mutants

To understand the substrate binding, release and catalytic mechanism, we introduced single mutations of Asp321, His61, Glu226 and Glu227 (**Figure 3**) and tested their activity using Michaelis-Menten kinetics. The pH conditions for forward and reverse reaction were selected based on the maximum rate constant obtained for each direction of the reaction. The catalytic rate constant, k_cat_, and Michaelis constant, K_m_, values were determined in the presence of excess of the second substrate for the forward reaction at pH 8.5 or the reverse reaction at pH 7.0 (**Figure 3a-c** and **Table S1-2**). Of note, K_m_ values reflect dynamic enrichment of enzyme molecules in the form of enzyme-substrate complex and thus represent apparent substrate affinity inherent to the steady state. For the forward reaction at pH 8.5, K_m_ values for ATP and Cr of WT uMTCK were determined to be 0.1 and 0.6 mM, respectively. K_m_ values for ADP and pCr for the reverse reaction at pH 7.0 were measured as 0.016 and 0.17 mM, respectively. The lower apparent affinity for Cr/pCr is consistent with our structural finding that suggested that Loop 2 can adapt closed conformation when ATP/ADP are bound in the active site. The catalytic rate constant was very similar between the forward and reverse reactions, ∼21 sec^-1^ and ∼40 sec^-1^, respectively.

**Figure 3.**
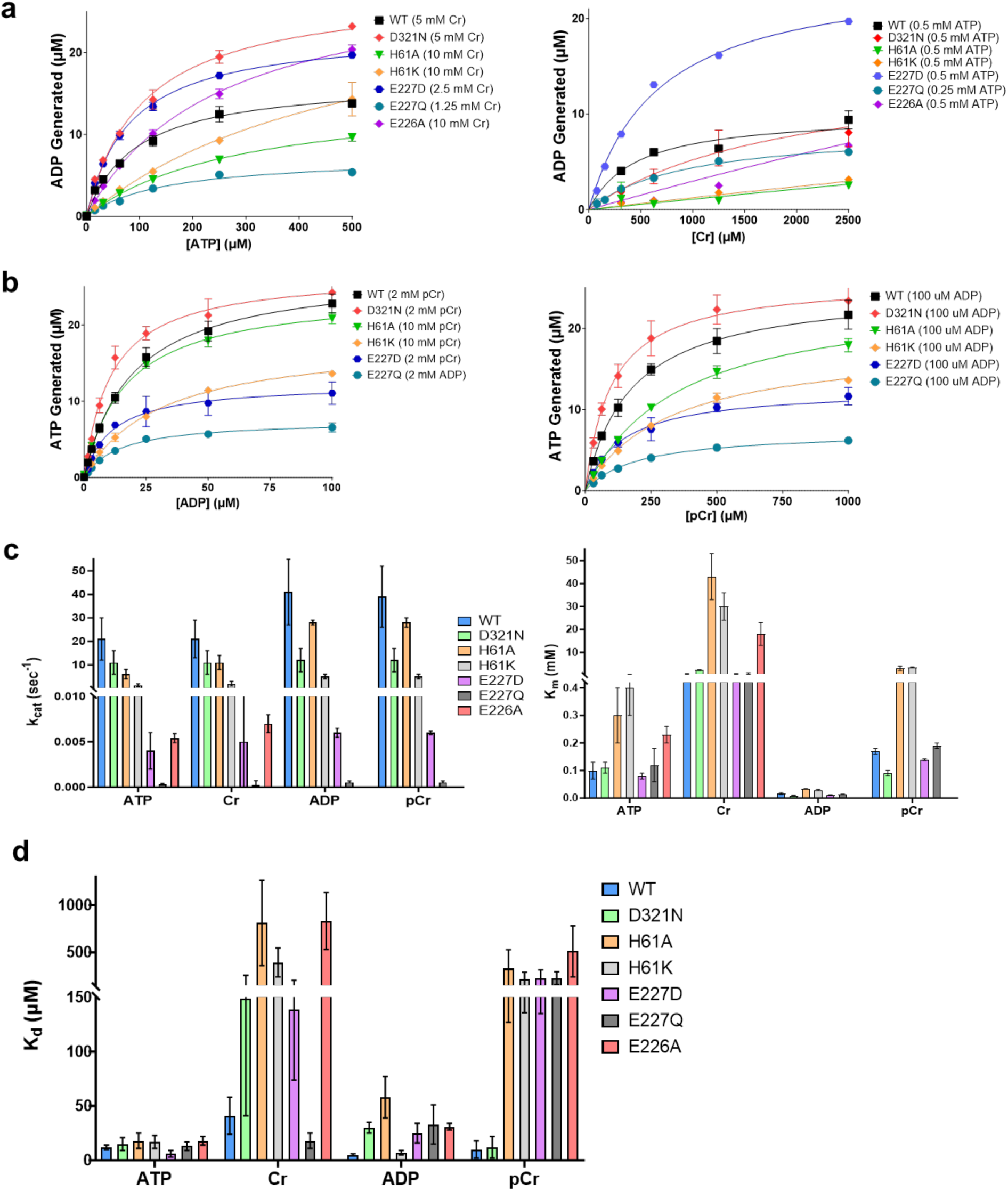
Representative curves from activity assays a) the k_cat_ and K_m_ values for ATP and Cr are measured in the forward reaction conditions at pH 8.5, and b) for ADP and pCr in the reverse reaction conditions at pH 7.0 c) Results of activity assays using at least three independent trials presented in bar graph format for comparison of k_cat_ and K_m_ values obtained for WT and mutant uMtCKs d) Results of binding assays from at least three independent trials displayed in bar graph format for comparison of K_d_ values for WT and mutant uMtCKs.

The catalytic rate constants of loop mutants, D321N, H61A and H61K, were comparable to the catalytic rate constant of WT uMtCK. The mutation of catalytic residue Glu227 to aspartate or glutamine resulted in a significant decrease in catalytic activity but did not abrogate activity completely. The mutation of Glu226 to alanine also exhibited a significant reduction in catalytic activity. Our finding indicates that this residue plays a key role in both substrate binding and catalytic activity.

While E227D and E227Q significantly decreased the catalytic activity of MtCK, the apparent affinity for all substrates remained the same. Based on our structural data, residue Glu226 is not in direct contact with the Cr substrate; however, E226A reduced Cr binding. Similarly, loop mutations H61A and H61K substantially decreased Cr and pCr binding, while D321N moderately increased K_m_ for Cr. The mutations of His61 revealed that the interaction between His61 and Asp321 is important to maintain Cr/pCr binding.

Indeed, mutation of His61 to alanine or lysine may have resulted in inability of Loop 2 to latch with Loop 1 to take the closed conformation to prevent the release of Cr from the active site. The replacement of Asp321 with asparagine maintains the residue’s ability to form a hydrogen bond with His61. However, the decreased affinity for Cr in the D321N mutant indicates Asp321 and His61 may form a stronger interaction such as a salt bridge that is lost upon mutation of Asp321 to an uncharged asparagine residue resulting in decreased affinity for Cr.

In contrast to the strong effect on Cr binding, the binding of ATP/ADP was not significantly affected by the mutations made to either of the loop residues. The slight increase in K_m_ values for ATP and ADP of H61A and H61K is potentially a reflection of using sub-saturating Cr concentrations due to the increase in K_m_ values for Cr. These findings indicate ATP/ADP binding is not dependent on the loops to latch together but rather on recognition of Mg^2+^-ATP/-ADP by interactions with the residues in the active site.

### Binding Cooperativity and Release of Substrates and Products

Next, we determined dissociation constants K_d_ values for each substrate and product using microscale thermophoresis under either forward or reverse reaction buffer conditions to compare with steady-state kinetic studies and to determine efficiency of product release from active site once the reaction is complete (**Figure 3d** and **Table 1**). The results of the binding assays for WT uMtCK confirmed our findings from the kinetics studies that show ATP/ADP is bound with a higher affinity than Cr/pCr in their respective reaction conditions. Furthermore, K_d_ values of ADP and pCr at pH 7.0 are lower than the respective K_d_ values obtained for ATP and Cr at pH 8.5. The binding affinities for the products of the forward reaction, pCr and ADP, at pH 8.5 were 10-fold lower than binding affinities for substrates Cr and ATP with WT uMtCK. Similarly, at pH 7.0, the binding affinities for substrates of reverse reaction pCr and ADP were higher than the binding affinities for products Cr and ATP. The increase of affinity towards substrates and loss of affinity for products in either forward or reverse reactions serves as a beneficial adaptation to increase the enzyme’s catalytic efficiency depending on the needs of cells.

**Table 1.**
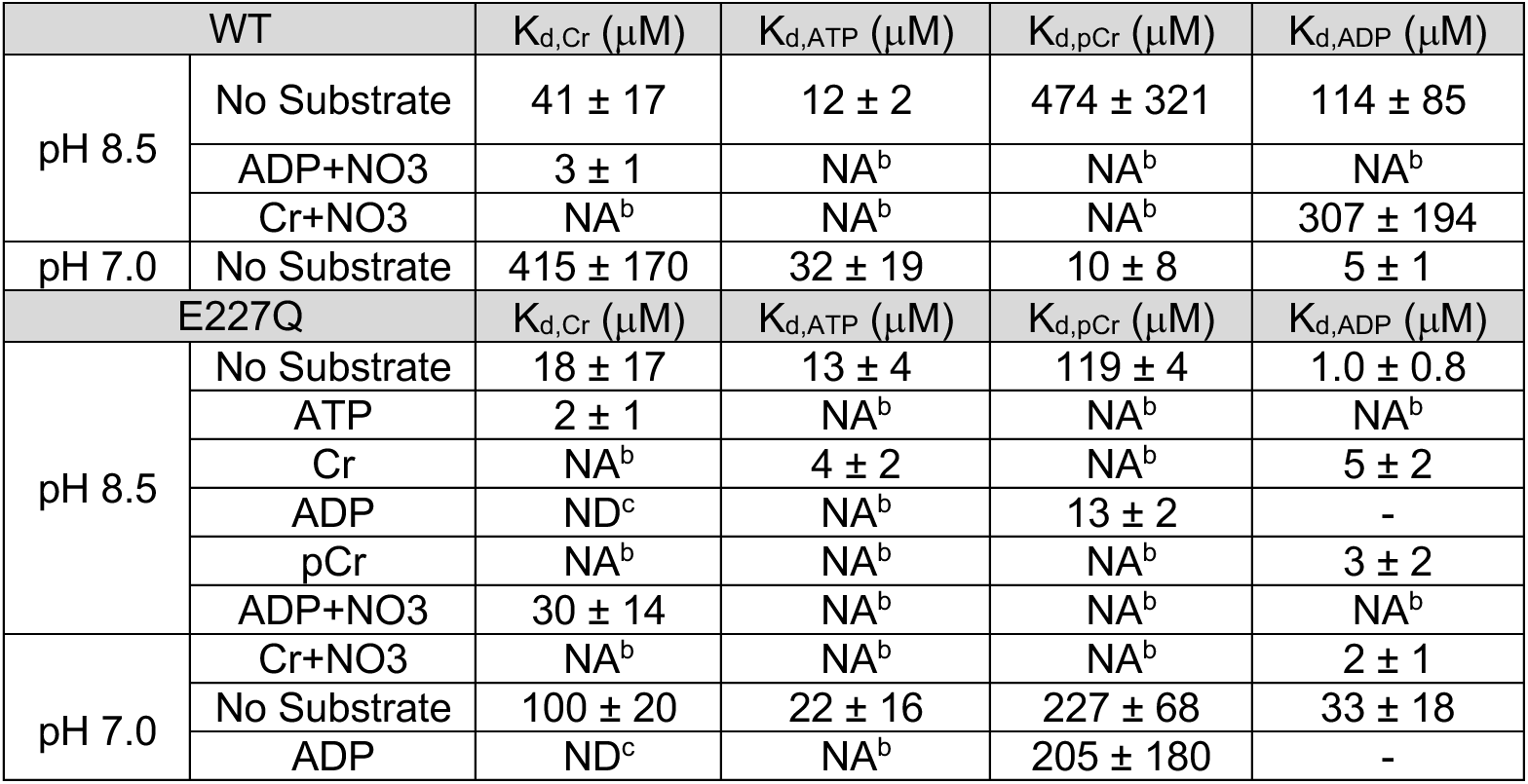
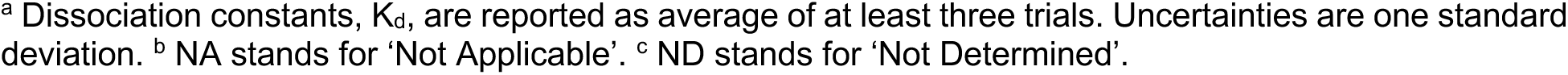
Binding data results for WT and E227Q uMtCK^a^.

| WT | | $K_{d,\text{Cr}}$ ($\mu\text{M}$) | $K_{d,\text{ATP}}$ ($\mu\text{M}$) | $K_{d,\text{pCr}}$ ($\mu\text{M}$) | $K_{d,\text{ADP}}$ ($\mu\text{M}$) |
| --- | --- | --- | --- | --- | --- |
| pH 8.5 | No Substrate | $41 \pm 17$ | $12 \pm 2$ | $474 \pm 321$ | $114 \pm 85$ |
| | ADP+$\text{NO}_3$ | $3 \pm 1$ | NA<sup>b</sup> | NA<sup>b</sup> | NA<sup>b</sup> |
| | Cr+$\text{NO}_3$ | NA<sup>b</sup> | NA<sup>b</sup> | NA<sup>b</sup> | $307 \pm 194$ |
| pH 7.0 | No Substrate | $415 \pm 170$ | $32 \pm 19$ | $10 \pm 8$ | $5 \pm 1$ |
| E227Q | | $K_{d,\text{Cr}}$ ($\mu\text{M}$) | $K_{d,\text{ATP}}$ ($\mu\text{M}$) | $K_{d,\text{pCr}}$ ($\mu\text{M}$) | $K_{d,\text{ADP}}$ ($\mu\text{M}$) |
| pH 8.5 | No Substrate | $18 \pm 17$ | $13 \pm 4$ | $119 \pm 4$ | $1.0 \pm 0.8$ |
| | ATP | $2 \pm 1$ | NA<sup>b</sup> | NA<sup>b</sup> | NA<sup>b</sup> |
| | Cr | NA<sup>b</sup> | $4 \pm 2$ | NA<sup>b</sup> | $5 \pm 2$ |
| | ADP | ND<sup>c</sup> | NA<sup>b</sup> | $13 \pm 2$ | - |
| | pCr | NA<sup>b</sup> | NA<sup>b</sup> | NA<sup>b</sup> | $3 \pm 2$ |
| | ADP+$\text{NO}_3$ | $30 \pm 14$ | NA<sup>b</sup> | NA<sup>b</sup> | NA<sup>b</sup> |
| pH 7.0 | Cr+$\text{NO}_3$ | NA<sup>b</sup> | NA<sup>b</sup> | NA<sup>b</sup> | $2 \pm 1$ |
| | No Substrate | $100 \pm 20$ | $22 \pm 16$ | $227 \pm 68$ | $33 \pm 18$ |
| | ADP | ND<sup>c</sup> | NA<sup>b</sup> | $205 \pm 180$ | - |

<sup>a</sup> Dissociation constants, $K_d$ , are reported as average of at least three trials. Uncertainties are one standard deviation. <sup>b</sup> NA stands for 'Not Applicable'. <sup>c</sup> ND stands for 'Not Determined'.

The evaluation of substrate binding with mutants has confirmed our observations from kinetic studies (**Table S1** and **S2**). Affinities for ATP or ADP were not affected by the mutations except for significantly reduced affinity for ADP with H61A mutant (**Figure 3d**). Binding of Cr or pCr were negatively impacted by mutations of Asp321, His61 and Glu226. As previously discussed, we believe the loss of affinity for Cr or pCr is due to loss of interactions that enable the loops to be latched together. In addition, mutation of the catalytic residue Glu227 significantly decreased the affinity of the enzyme for pCr.

To understand cooperativity between binding of substrates, we tested the binding affinity of Cr in the presence of ADP and NO_3_^-^ at pH 8.5. The results revealed Cr binds with a higher affinity once transition state mimics are bound in the active site. The binding of ADP was not affected by the presence of Cr and NO_3_^-^. To assess the cooperativity in substrate binding using natural substrates Cr and ATP without the interference of the reaction, we next obtained the binding affinities of Cr or ATP in the presence of each other using E227Q mutant. Both Cr and ATP showed cooperativity in binding in the presence of the other substrate. The increased affinity for one substrate in the presence of the other may be the result of stabilization of the loops upon binding of the first substrate that prevents the release of the second substrate as loops latch together. Interestingly, Cr or ADP binding in the presence of NO_3_^-^ representing TSA showed no cooperativity at pH 8.5 unlike our observation with WT enzyme.

In the conditions utilized in the reverse reaction (pH 7.0), the binding of pCr and ADP was impacted by mutation of catalytic residue Glu227 to glutamine resulting in a significant decrease in affinity compared to WT. In addition, pCr and ADP exhibited comparable binding affinity to Cr and ATP, and no cooperativity in binding. Interestingly, in the conditions used for the forward reaction (pH 8.5), E227Q has higher affinity for pCr and ADP than at pH 7.0, furthermore pCr shows cooperativity in binding in the presence of ADP. Inversely, Cr appears to have higher affinity to the E227Q mutant in both forward and reverse direction.

Based on the binding assays performed at two different pH conditions, modification of the catalytic residue proved to complicate the evaluation of substrate binding and their cooperativity. The modifications may have resulted in an altered reaction mechanism due to significantly reduced rate of reaction or changes in electrostatic environment in the active site upon mutation of the catalytic residue. Hence, we aimed to obtain structures of E227Q uMtCK in the presence of TSA or substrates/products to reveal detectable active site structure changes that could hint to possible changes in mechanism.

### TSA-MtCK_E227Q_ and Prod-MtCK_E227Q_ Structure Determination and Implication

Cryo-EM structures of E227Q mutant bound to either TSA complex (TSA-MtCK_E227Q_) or products ADP and pCr (Prod-MtCK_E227Q_) both at pH 8.0 and 7.0 were obtained to further our understanding of the reaction mechanism. The density maps obtained at pH 8.0 and 7.0 for both structures of TSA-MtCK_E227Q_ and Prod-MtCK_E227Q_ were identical, hence we are only presenting the structures obtained at pH 8.0 for simplicity. E227Q structure in complex with TSA showed a density for NO_3_^-^ ion that was absent in the structure of TSA-MtCK_WT_ (**Figure 4a**). Mutation of negatively charged glutamate to uncharged glutamine may have increased residence time of the negatively charged NO_3_^-^ ion in the active site. Surprisingly, Prod-MtCK_E227Q_ structure showed similar densities in the active site as the TSA-MtCK_E227Q_ structure (**Figure 4b**). ADP is bound in its binding pocket as expected, however the density by Cr/pCr binding site is too small to be phosphocreatine and lacks connected density for the phosphoryl group of pCr. Indeed, we suspect an extra density observed between ADP\ATP and Cr/pCr binding sites suggests pCr may have been converted to Cr and the extra density between the two sites may belong to a free phosphate group that is formed upon dephosphorylation of pCr by E227Q uMtCK. Previous studies on cCKs proposed that phosphoryl transfer proceeds through an S_N_2 mechanism. Our finding suggests that mutation of catalytic Glu227 residue to a glutamine may have modified the reaction mechanism to catalyze unproductive dephosphorylation of pCr that creates Cr and a free phosphate. In conclusion, the structures of E227Q uMtCK revealed that the electrostatic changes in the active site due to the replacement of the catalytic residue appear to alter the reaction mechanism and binding of substrates or TSA.

**Figure 4.**
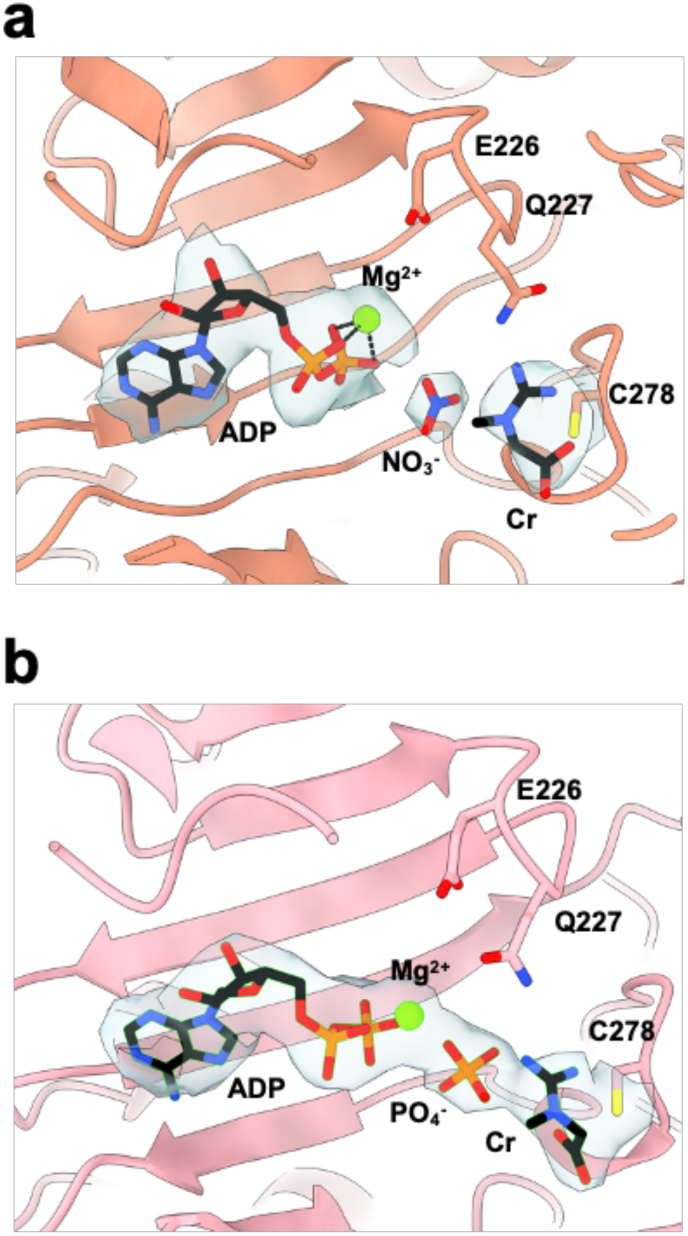
Structures of mutant E227Q uMtCK. a) The structure of TSA-MtCK_E227Q_ shows extra density between ADP and Cr that represent the NO_3_^-^ ion that mimics the transition state of phosphoryl transfer. b) The structure of Prod-MtCK_E227Q_ displays similar density between ADP and Cr that was modeled with PO_4_^-^.

### Inhibition mechanism of Covalent Inhibitor CKi

CKs role in different cancer types has been well-established, hence one of the main reasons for studying these enzymes to understand their role and functionality is to design selective and potent inhibitors that can target cancer cells. A covalent inhibitor, CKi, was described in a recently reported study. Its discovery leveraged the presence of a reactive active site cysteine residue by using chemoproteomics method and screening a cysteine-reactive small molecule library with CPT-MS^31^. The reactive cysteine residue of CKs corresponds to Cys278 in uMtCK that is located in the Cr binding site and is involved in the substrate binding alongside of Glu227 (**Figure 2d**). As part of the same study, covalent attachment of the inhibitor was verified with an X-ray structure of CKB (creatine kinase, brain-type) bound to the inhibitor confirming that it competes with substrate Cr for binding^31^. While the impact of CKi in UCSD-AML1 cells was examined in this study, its effect on different cancer types remains unknown. To understand the mechanism, selectivity and evaluate impact of CKi on breast cancer cells overexpressing MtCK, we sought out to test the inhibitor *in vitro* and in cellular assays using HER2+ breast cancer BT474 cell line.

Our *in vitro* and binding studies showed that CKi inhibits uMtCK activity after two-hour pre-incubation with an apparent IC_50_ value of 3.0 μM and binds with an apparent K_d_ of 13 μM (**Figure 5b-c**). Next, we evaluated commercially available analogs of CKi lacking the chloroacetamide group to assess the ability of the small molecule to bind in the active site without formation of a covalent link (**Figure 5a-c**). Surprisingly, the noncovalent analogs of CKi bind with similar affinity as CKi, however the activity assay with the analogs has not shown any inhibition effect on uMtCK. These observations confirmed that CKi inhibitor is non-selective pan-creatine kinase inhibitor, and its effects require the formation of the covalent bond with Cys278. We obtained a cryo-EM structure of uMtCK bound to CKi and ADP (CKi-MtCK_WT_) to understand how the presence of ADP/ATP affects CKi binding (**Figure 5d**). The previously obtained crystal structure of CKB with CKi alone had shown no conformational changes upon CKi binding where the loops were in open conformation. Our CKi-MtCK_WT_ structure showed a continuous extra density by Cys278 that indicated CKi is covalently attached at the Cr binding site with Cys278 similar to what is observed in the CKB structure and despite the ADP binding, both Loop 1 and 2 were found in open conformation. This finding indicates uMtCK is highly specific to Cr binding in the active site for the loop closures to occur and other small molecule binding with ADP/ATP is not as recognizable for loop closure and initiation of the reaction. In addition, the structure of CKi-MtCK_WT_ showed that the inhibitor makes a limited number of interactions with residues Thr54 and Gly68 in the active site. These results suggest that CKi might be guided into the active site by weak interaction with near-by residues but the inhibitory effect of CKi relies mainly on covalent link formation that abrogates Cr binding and catalytic reaction.

**Figure 5.**
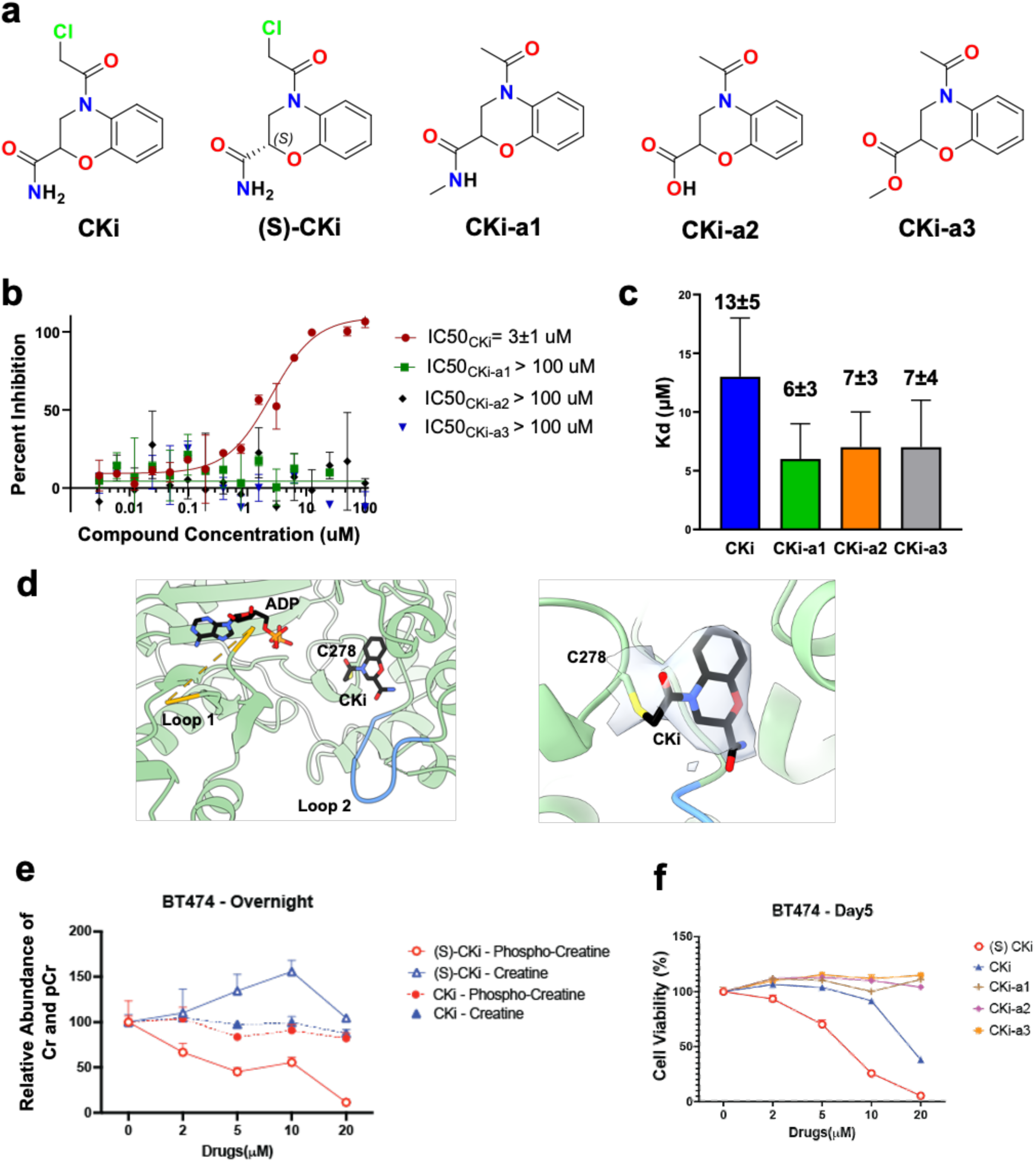
Analysis of covalent inhibitor CKi and its analogs a) Structures of cysteine-reactive small molecule CKi and its noncovalent analogs b) Representative dose-response curves for CKi and its analogs showed no inhibition of uMtCK with noncovalent analogs of CKi and inhibition with IC_50_ value of 2.5 μM with CKi obtained from at least three independent trials c) Dissociation constants of CKi and its analogs obtained from at least three independent trials are comparable, even though noncovalent analogs showed no inhibition of uMTCK d) In the CKi-MtCK_WT_ structure, CKi is bound in the Cr binding site while ADP is bound in the ATP/ADP binding site, however Loop 1 and 2 are in open conformations. The density around CKi and C278 residue is shown in the second panel. e) LC-MS analysis of creatine and phosphocreatine levels in BT474 cells treated with CKis for overnight. f) Proliferation assay results of BT474 cells treated with different CKi variants for 5 days.

We next sought to measure levels of the MtCK1 substrate and product (creatine and phosphocreatine) in breast cancer BT474 cells using liquid chromatography-mass spectrometry (LC-MS) and examine if racemic or more potent (S)-CKi treatment alter these intracellular levels. Overnight treatment of breast cancer BT474 cells with (S)-CKi decreases pCr levels in a dose-dependent manner, which is consistent with a reported study using leukemia cell lines (**Figure 5e**)^31^. However, the racemic CKi treatment did not reduce intracellular pCr levels even at 20 mM, indicating the importance of chirality in CKi derivatives to exhibit its inhibition effects on MtCK1. We further assessed the compounds’ efficacy against BT474 cell proliferation. After 5 days of treatment, BT474 cells exhibited varying responses to different CKi variants (**Figure 5f**). (S)-CKi was found to be the most potent inhibitor, achieving an IC_50_ value of 7.5 μM. The racemic CKi, while still effective, showed reduced potency with an IC_50_ value of 17.5 μM. Notably, other noncovalent analogs of CKi failed to inhibit BT474 cell proliferation within the same dosage range, supporting the *in vitro* results that indicate covalent bond formation by CKi plays a critical role in its inhibition effects.

Taken together, our findings suggest that CKi is not an effective inhibitor of uMtCK due to 1) its inability to bind with high affinity and mainly depend on reversible covalent bond formation and 2) lack of conformational changes in Loop 1 and 2 that can lock the inhibitor in position. Based on structural and biochemical data, we believe that allosteric inhibitors that can induce conformational changes and affect substrate/product binding indirectly would be a greater interest to target MtCK.

## Conclusion

We set out to investigate the key mechanistic features of ubiquitous mitochondrial creatine kinase using mutational, biochemical and structural analysis. Our findings indicated ATP binding and Loop 1 closure are independent of Cr binding, while Loop 2 closure mainly depends on Cr binding and is enabled by salt bridge formation between His61 and Asp321. Furthermore, the structures of catalytically rendered E227Q uMtCK showed that electrostatic changes to the active site can alter the mechanism of the reaction and substrate binding. Finally, we evaluated the inhibition mechanism of the only known up-to-date inhibitor, CKi, and revealed the covalent bond dependency of the inhibitor and the lack of conformational changes in the enzyme upon inhibitor binding even in the presence of substrate ADP. We showed that CKi is a non-selective inhibitor that inhibits both cytoplasmic and mitochondrial creatine kinases, thus is expected to indiscriminately shut down cellular processes and result in high overall cytotoxicity. We believe that identification of selective MtCK1 inhibitors with diverse mechanisms of action would provide a toolbox of chemical agents that would help establish pharmacologic approaches for targeting MtCK functions specific to cancer cells. Optimization of these newly identified inhibitors would rely on cryo-EM approach and would benefit from ability to obtain structures of uMtCK in complexes with small molecule ligands, as we readily demonstrated in this publication.

## Methods

A detailed methods and material section is described in the Supporting Information.

## Data Availability

The cryo-EM maps of MtCK are deposited to the Electron Microscopy databank with accession numbers of EMDB-44029, EMDB-44068, EMDB-44028, EMDB-44055, EMDB-44058 and EMDB-44069. The model coordinates of MtCK structures are deposited in Protein Data Bank with PBD codes of 9B05, 9B14, 9B04, 9B0T, 9B0U and 9B16. **Table S3** includes a summary of cryo-EM data collection, refinement and validation statistics.

## Acknowledgements and Funding Sources

This work was supported by NIH/NCI grant (R01 CA251910) to Eduard Sergienko and Taro Hitosugi. Cryo-EM data was collected at Cryo-EM facility at Sanford Burnham Prebys Medical Discovery Institute supported by NIH/NCI grants (S10 OD026926 and P30 CA030199). MtCK1 protein production and MST were carried out at the Protein Production and Analysis Facility supported by NIH/NCI grant (P30 CA030199). Merve Demir was supported by Conrad Prebys Foundation award number CPF #651.

## Notes

### Competing Interest Statement

The authors have declared no competing interest.

### Summary of Updates

The yellow highlight on page 24, paragraph 2 was removed.

