## Supporting Information for "Structural Basis for Substrate Binding, Catalysis and Inhibition of Breast Cancer Target Mitochondrial Creatine Kinase by Covalent Inhibitor via Cryo-EM"

### Methods and Materials

#### Protein Production

The pet15b vector containing human uMtCK gene was synthesized by GenScript. Single point mutations of uMtCK were generated using appropriate primers ordered from Eton Biosciences and verified by sequencing with Primordium. Rosetta 2 (DE3) cell lines were transformed with pet15b vector overexpressing human uMtCK or its mutants with N-terminal His-tag with TEV cleavage site and C-terminal FLAG-tag. Protein was overexpressed with 0.5 mM IPTG overnight at 15 C. Cell pellet was resuspended in lysis buffer (50 mM Tris-HCl pH 8.0, 200 mM NaCl and 5% glycerol) supplemented with protease inhibitor, DNase1 and 10 mM $MgCl_2$ and lysed by sonication. Protein was purified using HisPur Co-NTA beads by gravity flow which is followed by size exclusion column HiLoad 26/600 Superdex 200 that showed multiple peaks for octamer, dimer and monomer of uMtCK. The peak for octameric protein in a final buffer condition of 50 mM Tris-HCl pH 8.0 and 200 mM NaCl was collected, and aliquots were stored at –80 C until use for cryo-EM or biochemical assays.

#### Activity Assay

The activity of human uMtCK and its mutants were tested by performing a time-dependent activity assay with serial dilution of proteins to determine the optimal reaction conditions such as incubation time and enzyme concentration. A cross-titration of substrates in optimal reaction conditions for each protein was carried out to measure the catalytic rate constant, $k_{cat}$, and Michaelis constant, $K_m$, in 1536 or 384 plate format. The values for ATP/ADP were determined in the presence of excess Cr, while values for Cr/pCr were determined in the presence of the other substrate. The final assay buffer conditions were composed of 25 mM Tris-HCl pH 8.5 for forward and pH 7.0 for reverse reaction, 7 mM $MgCl_2$, 250 mM sucrose, 2 mM DTT and 0.005% Tween 20. The ADP or ATP product formation was measured using ADP-Glo or Kinase-Glo kit by Promega and Pherastar FS microplate reader to measure luminescence output. Data were analyzed using GraphPad Prism and fitting the curves to the Michaelis-Menten equation. The small molecule inhibitor of racemic CKi and its analogs were purchased from Enamine and (S)-CKi from ProbeChem. Serially diluted compounds were pre-incubated with 0.45 nM WT uMtCK in 25 mM Tris-HCl pH 8.5, 7 mM $MgCl_2$, 250 mM sucrose, 2 mM DTT and 0.005% Tween 20 assay buffer for 2 hours at room temperature. The final product formation was measured after 30 min incubation of compound-protein mixture with 0.25 mM ATP and 0.8 mM Cr at room temperature using ADP-Glo kit from Promega.

**Microscale Thermophoresis**

uMtCK and its mutants were buffer exchanged to 50mM $PO_4^{3-}$ pH 8.0 and 150 mM NaCl for conjugation reaction. Protein was incubated with Dylight 650 NHS ester for 1 hour at room temperature on rocker at dark. Conjugation reaction is quenched with 10 mM glycine and the reaction mixture is dialyzed to the storage buffer (50 mM Tris-HCl pH 8.0 and 150 mM NaCl). Aliquots of labeled protein were stored at –80 C in the dark.

For the binding assay, substrates were serially diluted using assay buffer (50 mM Tris-HCl pH 8.5 for forward and pH 7.0 for reverse reaction, 5 mM $MgCl_2$, 100 mM NaCl and 0.05% Tween 20) and mixed with equal volume of protein diluted in assay buffer to a concentration of 50 nM before mixing with substrate. The samples were run on NanoTemper Monolith NT.115 MicroScale Thermophoresis instrument to measure the amount of unbound and bound protein across varying concentration substrates and data was fit into a single binding isotherm to extract the $K_d$ value. For testing of CKi that is dissolved in DMSO, the assay buffer was adjusted with 5% final DMSO concentration to account for the DMSO carried over from the initial dilution of the inhibitor. The binding affinity of compounds tested in the presence of substrate or TSA included 1 mM ATP/ADP, 10 mM Cr/pCr or 50 mM $NO_3^-$.

**Cryo-EM**

Cryo-EM sample preparation and data collection was performed in the Sanford Burnham Prebys Cryo-EM Core facility. 4ul of 0.5-1 mg/mL of uMtCK protein supplemented with 50 mM $MgCl_2$ was applied onto a Quantifoil holey carbon grid (R2/2, 300mesh, copper) (glow-discharged in a Pelco easi-glow at 15 mA for 25 s) in a Vitrobot MarkIV, blotted for 10 s at 4 °C and 100% humidity (blot force 0) and subsequently plunge-frozen in liquid ethane. For samples with substrates, protein was preincubated with 10 mM Cr, 1 mM ADP and 50 mM $NO_3$ for 15 minutes. 1 mg/mL MtCK with 50 mM $MgCl_2$ was preincubated with 0.5 mM CKi and 1 mM ADP for 1.5 hours to obtain the CKi-$MtCK_{WT}$ structure.

The grids were imaged in a Titan Krios microscope (Thermo Fischer) at 300kV. Cryo-EM data acquisition was performed using SerialEM[1] by image shift (four shots per hole) with beam-tilt compensation. Movies with 20 frames each were recorded using a Gatan K3 detector in super resolution mode with a dose rate of around 20 electrons per pixel per second and with a total electron dose of around 34 e-/A2, at a pixel size of 1.06A per physical pixel. All micrographs were collected with a defocus ranging between 0.9 to 2.1 μm.

Data processing was carried out in Cryosparc 3.3.2[2]. The collected movies were subjected to motion-correction and dose-weighting of frames by Patch motion correction using default parameters. The CTF parameters were estimated by Patch CTF estimation and binned by 2. Images with ice contamination, failed CTF estimation or low resolution CTF fit values were discarded. For MtCK particle picking was done initially with the blob picker function in cryoSPARC using a round blob with a diameter of 135A to 155A and extracted using a box size of 256 pixels and binned by 4. The binned particles were cleaned by 2D classification. An Ab-Initio reconstruction with a D4 symmetry enforced was generated from the unbinned particles followed by a 3D reconstruction using Non-uniform Refinement. The resulting 3D map was used to prepare 40 equally-spaced 2D templates, which were then used as templates for template-based picking using a mask diameter of 145A. Particles were again extracted using a box size of 256 pixels, binned by 4 and junk particles were removed by 2D classification. Unbinned particles were used to generate an Ab-Initio reconstruction with D4 symmetry enforced, followed by a 3D reconstruction using Non-uniform Refinement. To confirm the D4 symmetry an additional Ab-Initio reconstruction without D4 symmetry enforced was generated, followed by a 3D reconstruction without D4 symmetry enforced.

The crystal structure 1QK1 was used as a starting model and was refined using an iterative process of manual adjustments and refinement (with Torsion, Planar Peptide, Trans Peptide, and Ramachandran restraints turned on) in Coot v.0.9.8.1[3,4] and subsequent refinement with phenix.real_space_refine in

Phenix v.1.20[5,6]. The final model was validated using MolProbity and Phenix. The resulting cryo-EM maps and models were visualized using UCSF Chimera[7].

**Cell Line Preparation**

Breast cancer BT474 cells were cultured in phenol red-free Dulbecco's Modified Eagle Medium (DMEM)/Ham's F12 medium supplemented with 10% fetal bovine serum (FBS), 100 U/mL penicillin G, and 100 µg/mL streptomycin (P/S). All the cells were cultured in 37 °C humidified incubators with 5% CO2.

**Metabolite Extraction and LC-MS analysis**

BT474 cells ($0.6 \times 10^6$ cells) treated with drugs for overnight were collected and washed by PBS. Intracellular metabolites were extracted with 250 µL of 10% TCA containing 1 µg D3 creatine. All samples were sonicated for 5 seconds and then centrifuged at 11,500 × g for 2 min at 4 °C. The supernatant was transferred to a new 1.5 mL tube. To remove the TCA, a fresh organic solvent mixture of trichlorotrifluoroethane (TCTFE):trioctylamine (3:1 ratio) was added to the supernatant at a ratio of 2:1 (organic solvent:TCA). The mixtures were vortexed for 15 seconds and allowed to separate at room temperature for 5 minutes. The aqueous layer was removed with a plastic pipette and transferred to a new 1.5 mL Eppendorf tube. The pH of each sample was adjusted to 8.0 using 2 µL of 1M Tris. The pH-adjusted samples were transferred into autosampler vials, and a 10 µL sample was injected for LC-MS analysis. Chromatographic separation was performed using an Agilent InfinityLab Poroshell 120 EC-C18 analytical column (3.0 × 50 mm, 2.7-micron) at 40 °C. The elution gradient consisted of LCMS-grade water with 0.1% formic acid (A) and 100% methanol (B) at a flow rate of 0.3 ml/min for a total run time of 30 min. The elution began at 100% A and was held for 0.01 min. Then, B was increased to 0.5% and held for 7.99 min. At a total time of 8 min, B was increased to 80% and held for 2 min. Subsequently, B was decreased to 0.5% and held for 0.01 min. Finally, at a total time of 15.01 min, B returned to 0%, returning to the initial conditions, and the run finished with 15 min at 100% A. The auto-sampler temperature was maintained at 4 °C. The retention times of the peaks were confirmed using freshly prepared pure standard compounds in LCMS-grade water. The relative abundance of peaks compared to their standards was collected from the Data Analysis of Agilent OpenLab Control Panel and analyzed using GraphPad Prism 9 version 9.5.1.

**Cell Proliferation Assay**

Cell proliferation was conducted using the CyQUANT Cell Proliferation Assay Kit (Thermo Fisher Scientific). Cells were seeded in a 96-well plate and treated with the creatine kinase inhibitors (CKis), including (S)-CKi, racemic CKi, CKi-a1, CKi-a2, and CKi-a3, at the indicated concentration. The cells were allowed to grow for 5 days under same culture conditions. After 5 days incubation, 50 μL of the detection reagent was added to each well, and the plate was incubated for 60 minutes at 37 °C. The fluorescence intensity was measured using a microplate reader with excitation and emission wavelengths set at 508 nm and 527 nm, respectively. The fluorescence signal was detected from the bottom of the well. The cell viability data obtained from the CyQUANT Cell Proliferation Assay were analyzed using GraphPad Prism 9 version 9.5.1.

**Table S1.** Kinetic and binding data results for uMtCK and its mutants at pH 8.5 for forward reaction conditions[a]

| **uMtCK** | $k_{cat,ATP}$ (sec$^{-1}$) | $K_{m,ATP}$ (mM) | $K_{d,ATP}$ (μM) | $k_{cat,Cr}$ (sec$^{-1}$) | $K_{m,Cr}$ (mM) | $K_{d,Cr}$ (μM) |
|---|---|---|---|---|---|---|
| WT | 21±9 | 0.10±0.03 | 12±2 | 21±8 | 0.6±0.1 | 41±17 |
| D321N | 11±5 | 0.11±0.02 | 15±6 | 11±5 | 2.2±0.2 | 149±108 |
| H61A | 6±2 | 0.3±0.1 | 18±7 | 11±3 | 43±10 | 812±451 |
| H61K | 1.3±0.6 | 0.4±0.1 | 17±6 | 2±1 | 30±6 | 394±153 |
| E227D | 0.004±0.002 | 0.08±0.01 | 6±3 | 0.005±0.002 | 0.6±0.1 | 139±65 |
| E227Q | 0.0003±0.0001 | 0.12±0.06 | 13±4 | 0.00022±0.00005 | 0.8±0.3 | 18±7 |
| E226A | 0.0054+0.0005 | 0.23±0.03 | 18±4 | 0.007±0.001 | 18±5 | 833±302 |

[a] Kinetic and binding parameter are reported as average of at least three trials. Uncertainties are one standard deviation.

**Table S2.** Kinetic and binding data results for uMtCK and its mutants at pH 7.0 for reverse reaction conditions[a]

| **uMtCK** | $k_{cat,ADP}$ (sec$^{-1}$) | $K_{m,ADP}$ (mM) | $K_{d,ADP}$ (μM) | $k_{cat,pCr}$ (sec$^{-1}$) | $K_{m,pCr}$ (mM) | $K_{d,pCr}$ (μM) |
|---|---|---|---|---|---|---|
| WT | 41±14 | 0.016±0.03 | 5±1 | 39±13 | 0.17±0.01 | 10±8 |
| D321N | 12±5 | 0.009±0.001 | 30±5 | 12±5 | 0.09±0.01 | 12±10 |
| H61A | 28±1 | 0.033±0.001 | 58±19 | 28±2 | 3±1 | 328±201 |
| H61K | 5±1 | 0.029±0.003 | 7±2 | 5±1 | 3.4±0.2 | 213±77 |
| E227D | 0.006±0.0005 | 0.0114±0.0005 | 25±9 | 0.006±0.0002 | 0.139±0.003 | 225±90 |
| E227Q | 0.0005±0.0002 | 0.013±0.001 | 33±18 | 0.0005±0.0002 | 0.19±0.01 | 227±68 |
| E226A | <0.0001 | ND | 31±3 | <0.0001 | ND | 511±270 |

[a] Kinetic and binding parameter are reported as average of at least three trials. Uncertainties are one standard deviation.

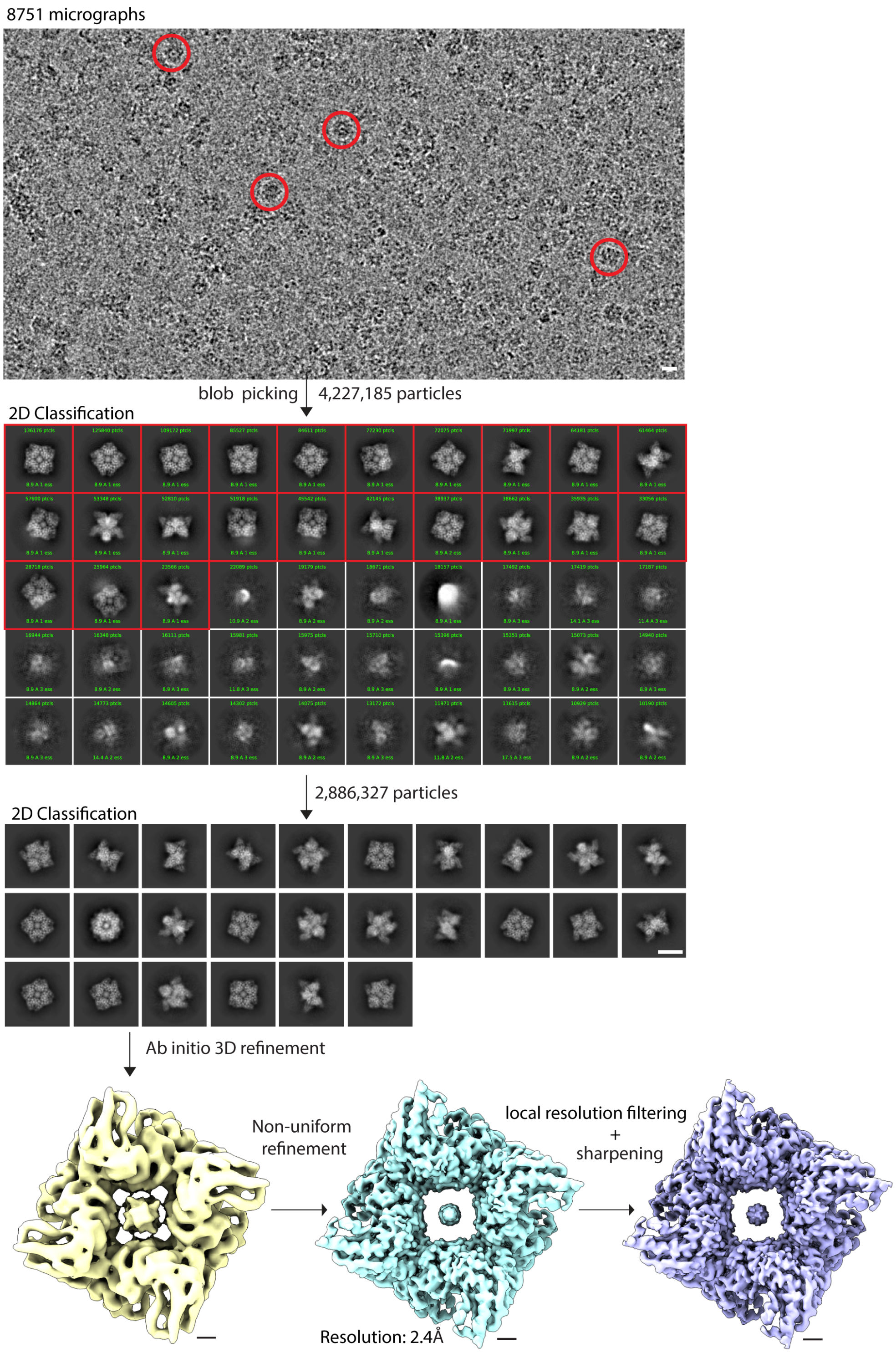


**Figure S1** Flowchart of cryo-EM data processing of MtCK using Cryosparc**.** On 8751 micrographs 4,227,185 particles were picked using the blob picker with a diameter range of 135Å to 155Å. After 3 rounds of 2D classification to remove junk and lower quality particles 2,886,327 particles remained and were subsequently used in the ab-initio reconstruction and Non-uniform refinement.

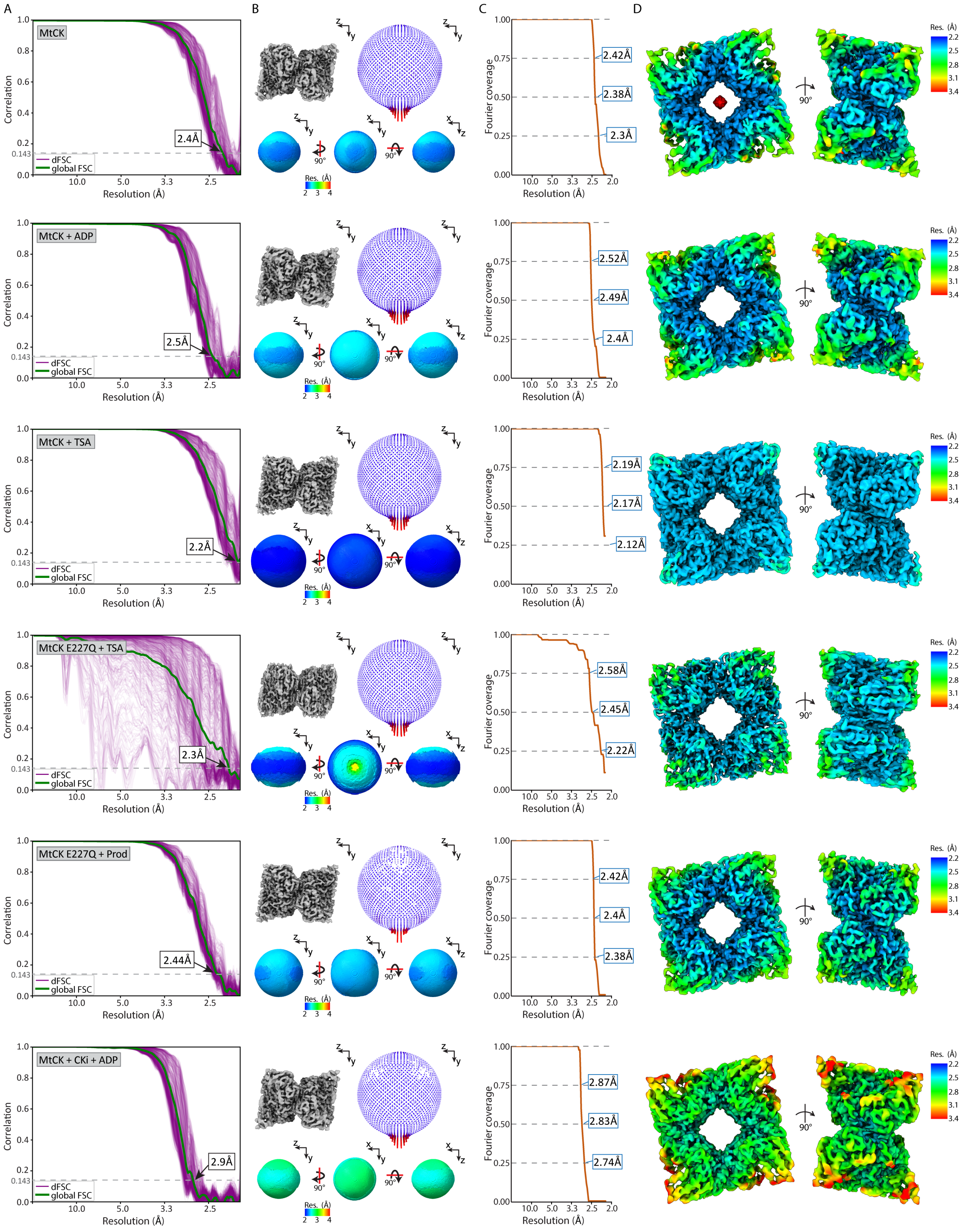


**Figure S2** Cryo-EM map statistics for Apo-, ADP-, and TSA-MtCK, TSA-$MtCK_{E227Q}$, Prod-$MtCK_{E227Q}$ and CKi-MtCK **A** Fourier shell correlation (FSC) and 1D directional FSC (dFSC) plots. **B** Visualizations of 3D dFSC. **C** Plot showing coverage of Fourier space. **D** Plot showing distribution of viewing angles from final refined dataset. **E** Final EM maps, colored by local resolution as estimated by Cryosparc.

**Table S3.** Cryo-EM data collection, refinement and validation statistics

| | Apo-MtCK EMDB-44029 PDB ID: 9B05 | TSA-MtCK$_{WT}$ EMDB-44068 PDB ID: 9B14 | ADP-MtCK$_{WT}$ EMDB-44028 PDB ID: 9B04 | TSA-MtCK$_{E227Q}$ EMDB-44055 PDB ID: 9B0T | Prod-MtCK$_{E227Q}$ EMDB-44058 PDB ID: 9B0U | CKi-MtCK$_{WT}$ EMDB-44069 PDB ID: 9B16 |
|---|---|---|---|---|---|---|
| **Data collection and processing** | | | | | | |
| Magnification | 47,170 | 47,170 | 47,170 | 47,170 | 47,170 | 47,170 |
| Voltage (kV) | 300 | 300 | 300 | 300 | 300 | 300 |
| Electron exposure (e–/Å$^2$) | 34 | 34 | 34 | 34 | 34 | 34 |
| Defocus range (μm) | 0.9-2.1 | 0.9-2.1 | 0.9-2.1 | 0.9-2.1 | 0.9-2.1 | 0.9-2.1 |
| Pixel size (Å) | 1.064 | 1.064 | 1.064 | 1.064 | 1.064 | 1.064 |
| Symmetry imposed | D4 | D4 | D4 | D4 | D4 | D4 |
| Initial particle images (no.) | 4,227,185 | 3,577,534 | 3,718,445 | 3,851,095 | 3,233,133 | 2,982,025 |
| Final particle images (no.) | 2,886,327 | 3,132,165 | 1,732,931 | 1,053,188 | 476,509 | 1,291,884 |
| Map resolution (Å)<br>FSC threshold | 2.40<br>0.143 | 2.20<br>0.143 | 2.52<br>0.143 | 2.30<br>0.143 | 2.44<br>0.143 | 2.89<br>0.143 |
| Map resolution range (Å) | 2.383 – 6.999 | 2.432 – 19.686 | 2.383 – 7.532 | 2.432 – 31.695 | 2.383 – 32.731 | 2.415 – 8.557 |
| **Refinement** | | | | | | |
| Initial model used (PDB ID) | 1QK1 | 1QK1 | 1QK1 | 1QK1 | 1QK1 | 1QK1 |
| Model composition<br>Protein residues<br>Ligands | 2720<br>0 | 2944<br>24 | 2944<br>16 | 2872<br>32 | 2944<br>32 | 2864<br>16 |
| R.m.s. deviations<br>Bond lengths (Å)<br>Bond angles (°) | 0.002<br>0.587 | 0.003<br>0.705 | 0.002<br>0.582 | 0.003<br>0.620 | 0.003<br>0.631 | 0.003<br>0.589 |
| Validation<br>MolProbity score<br>Clashscore<br>Poor rotamers (%) | 1.52<br>9.55<br>0.55 | 1.83<br>7.70<br>0.61 | 1.62<br>6.27<br>0.89 | 1.93<br>10.25<br>1.01 | 1.77<br>6.24<br>0.75 | 1.78<br>7.78<br>0.51 |
| Ramachandran<br>Favored (%)<br>Allowed (%)<br>Ouliers (%) | 97.99<br>2.01<br>0 | 93.82<br>5.77<br>0.41 | 95.97<br>4.03<br>0 | 94.12<br>5.88<br>0 | 93.44<br>6.28<br>0.27 | 94.85<br>5.15<br>0 |
